# Antioxidant modulation of stress behavior depends on stress coping style: a role for N-acetylcysteine amide

**DOI:** 10.64898/2026.08.10.741552

**Authors:** Ryan Y. Wong, Brooklynn K. Schmidt, Cherylynn R. Gibson, Peter D. Dijkstra

**Affiliations:** University of Nebraska at Omaha, Department of Biology, Omaha, NE USA; University of Nebraska at Omaha, Department of Psychology, Omaha, NE USA; Central Michigan University Biology Department, Mount Pleasant, MI USA

**Keywords:** stress, stress coping style, *N*-acetylcysteine amide, antioxidant, oxidative stress

## Abstract

Animals experience stressors in a variety of contexts that result in activation of neuroendocrine and cellular stress responses. Release of stress hormones can disrupt or restore redox homeostasis, and the resulting changes in oxidative states, physiology and behavior vary by an individual’s stress coping style. However, oxidative stress can also directly modulate neuroendocrine stress signaling. To what extent individual differences in brain antioxidant levels alter behavioral stress levels is not well understood. The present study investigated how *N*-acetylcysteine amide (NACA), an antioxidant and glutamate-modulating compound, regulates stress behavior across zebrafish (*Danio rerio*) with different stress coping styles (proactive, reactive). Following 24-hour exposure to NACA or control conditions, we quantified individual and composite stress behaviors using a Light–Dark Test (LDT). As expected, both proactive fish and NACA-treated fish showed significantly lower stress behaviors compared to reactive and control animals, respectively. Notably, stress-reducing effects of NACA were only seen in those with a reactive stress coping style. Overall, our data suggest that antioxidant mechanisms (e.g., glutathione system) may be key in facilitating the distinct behavioral and physiological responses to stressors that characterize alternative stress coping styles. The results underscore how individual differences in stress coping style and redox state can influence behavioral responses to stress.

## 1. Introduction

Animals are frequently exposed to stressors and display behavioral, physiological, and cellular responses to successfully overcome them. When exposed to a stressor, the neuroendocrine and autonomic stress axes activate to allow for fast modulation of physiology and behavior. While stress hormones directly influence neural activity in the brain through altering protein activity, oxygen, and energy utilization in the cell (Harris, 2020; McEwen et al., 2015; McEwen et al., 2012; Russo et al., 2012; Sanacora et al., 2022), the effects and resulting behavioral responses vary between individuals (Coppens et al., 2010; de Boer et al., 2017; Gilmour et al., 2025; Koolhaas et al., 2010; Overli et al., 2007). Factors contributing to variation in the magnitude and breadth of the effects include the individual’s intracellular biochemical state (e.g., redox homeostasis) and stress coping style (Baker et al., 2017; Bhattacharya et al., 2024; Coppens et al., 2010; Koolhaas et al., 2010; Sies et al., 2024).

Cellular redox homeostasis is achieved when there is a balance between prooxidant and antioxidant molecules. Excessive prooxidants (e.g., reactive oxygen species) compared to antioxidants leads to oxidative stress, which result in molecular damage and altered cellular signaling (Sies et al., 2024). In the nervous system, oxidative stress can impair neuronal function, disrupt neurotransmitter balance, and interfere with neuroendocrine stress signaling (Behrouzi et al., 2022; Bhattacharya et al., 2024; Cobley et al., 2018; Houldsworth, 2024; Salim, 2017; Sies et al., 2024). A variety of biotic and abiotic stressors can disrupt redox homeostasis leading to altered oxidative stress and antioxidant profiles (Bouayed et al., 2009; Costantini et al., 2011; Culbert et al., 2022; Dijkstra et al., 2024; Rammal et al., 2008; Sunday-Jimmy et al., 2026).

Antioxidants (e.g. glutathione) can mitigate the effects of neuroendocrine stress response and its behavioral manifestations by directly altering neurotransmission in the brain and reducing oxidative stress (Fedoce et al., 2018; Halliwell, 2024; Mocelin et al., 2015; Pamplona & Costantini, 2011; Prevatto et al., 2017). One antioxidant, N-acetylcysteine (NAC) and its amide derivative, N-acetylcysteine amide (NACA), has indirect antioxidant effects through being a precursor for glutathione and can directly influence neurotransmitter signaling (Bradlow et al., 2022; Pedre et al., 2021; Raghu et al., 2021; Spilere et al., 2025). Furthermore, administering NAC or NACA reduced stress and anxiety-like behaviors across a variety of animals (Dean et al., 2011; Marcon et al., 2019; Mocelin et al., 2015; Mocelin et al., 2018; Mocelin et al., 2019; Reis et al., 2020; Santos et al., 2017; Shariatmadari et al., 2026). Despite studies showing that altering oxidative status and antioxidant capacities (via redox signaling) is sufficient to change behavioral and hormonal stress states, the role of individual differences in shaping the link between oxidative state and stress coping is not well studied (Bouayed et al., 2009; Distler & Palmer, 2012; Hovatta et al., 2005; Mocelin et al., 2015; Mocelin et al., 2019; Prevatto et al., 2017; Reis et al., 2020; Salahinejad et al., 2021; Salim et al., 2010; Salim et al., 2011; Sies & Jones, 2020).

Individual differences in behavioral and physiological responses to a stress are seen across many animals (Baker et al., 2017; Koolhaas et al., 2010; Overli et al., 2007; Sih et al., 2004). A stress coping style is marked by a correlated suite of behavioral and physiological responses that are consistent across time and context, and ultimately facilitate an effective recovery from a stressor. Two qualitatively different stress coping styles seen in many vertebrates are the proactive and reactive stress coping styles (Baker et al., 2017; de Boer et al., 2017; Koolhaas et al., 2010; Overli et al., 2007). Individuals with a proactive style are characterized by being risk-prone, relying on a feed-forward memory process, possessing low behavioral flexibility, and exhibiting a relatively low glucocorticoid stress response. On the other end of the spectrum, individuals with a reactive style have diametrically opposing responses (Baker et al., 2017; Coppens et al., 2010; Koolhaas et al., 2010; Overli et al., 2007; Sih et al., 2004). Both coping styles are natural adaptive responses to challenges in the environment and may be maintained in a population due to differential fitness in variable environments (Dochtermann & Dingemanse, 2013; Sih et al., 2004). These stress coping styles have a genetic basis and consistently bias an individual’s behavioral and physiological responses across contexts and time (Baker et al., 2017; Koolhaas et al., 2010; Overli et al., 2007; Sih et al., 2004). Only a few studies have investigated the role of stress coping styles on oxidative stress and antioxidant capacities (Arnold et al., 2014; Coccaro et al., 2016; Costantini et al., 2008; Filiou et al., 2014; Herborn et al., 2011; Pamplona & Costantini, 2011; Sunday-Jimmy et al., 2026). Animals with a reactive-like coping style have elevated biomarker levels of oxidative stress and antioxidant activity at baseline or in response to an acute stressor, but this relationship can be complex and tissue specific (Arnold et al., 2014; Coccaro et al., 2016; Costantini et al., 2008; Filiou et al., 2014; Herborn et al., 2011; Pamplona & Costantini, 2011). Studies to date have focused on investigating changes in oxidative states between the stress coping styles in response to neuroendocrine stress but have not attempted to examine the effects of altering oxidative states on the stress response.

In this study we investigate effects of increasing an antioxidant (N-acetylcysteine amide, NACA) on individual differences in behavioral stress levels. We test the hypothesis that NACA exposure would reduce stress behaviors in a coping-style–dependent manner. After exposure to either NACA or vehicle control, we quantified behavioral stress level using a scototaxis assay (light-dark test) (Maximino et al., 2010). By linking antioxidant modulation with intrinsic stress coping styles, this study helps clarify sources of individual variability in stress-related behavior and informs the interpretation of antioxidant-based approaches to stress research.

## 2. Methods

### 2.1 Animals and Housing

We used two selectively bred lines of zebrafish (*Danio rerio*) that display, on average, the proactive or reactive stress coping styles. These lines originated from a population of wild caught zebrafish near the village of Giaghata, India and were generated and maintained in the laboratory by artificial selection for amount of movement in response to a novelty stressor in the open field test (Wong et al., 2012). Subsequent studies showed that these lines reliably differ in behavioral stress responses across different behavioral stress assays, time (weeks and generations), and novelty stressor-induced cortisol release rates that are consistent with the proactive and reactive stress coping styles (Baker et al., 2018; Johnson et al., 2020; Wong et al., 2019; Wong et al., 2012). Furthermore, these lines differ in cognition, morphology, whole-brain transcriptome profiles, functional brain network activity, whole-brain oxidative stress and antioxidant biomarker levels, and response to anxiogenic and anxiolytic compounds (Ayayee & Wong, 2024; Baker et al., 2017; Baker & Wong, 2019, 2021; Corcoran, Rushlau, et al., 2025; Corcoran, Storks, et al., 2025; Goodman & Wong, 2020; Kern et al., 2016; Klucas & Wong, 2026; Sunday-Jimmy et al., 2026; Wong et al., 2015; Wong et al., 2014; Wong et al., 2013). In this study fish were 11 months old and had undergone 14 generations of selective breeding. Prior to experiment, fish were maintained in 40L mixed-sex tanks on custom-built recirculating water system with solid and biological filtration. Water parameters consisted of 27 ± 1°C temperatures, pH of 7.2, conductivity of 700 µS, and fish were exposed to a 14:10 hour light/dark cycle. Fish were fed twice daily with Tetramin Tropical commercial flake food.

### 2.2 NACA treatment and behavioral stress assay

We randomly assigned fish to four experimental groups: NACA-treated proactive fish (n=3 females; n=9 males), control-treated (water) proactive fish (n=5 females; n=7 males), NACA-treated reactive fish (n=6 females; n=6 males), and control reactive fish (n=6 females; n=6 males). Sex was determined by visualization of ovaries or testes on dissection. In the current study we followed modified NACA treatment procedures from a previously published study using NACA in adult zebrafish (Reis et al., 2020). NACA is rapidly converted to NAC after uptake, and has enhanced ability to cross the blood-brain barrier and antioxidant properties compared to NAC (He et al., 2020; Reis et al., 2020). Groups of 4 fish were submersed in water containing either 1 mg/L NACA (Millipore Sigma A0737) or solvent (water) for 24 hours.

Following the 24-hour exposure period, we quantified behavioral stress levels using the light dark test (LDT) following a previous protocol (Wong et al., 2012). In brief, a 30 × 30 × 10 cm tank filled with 4L of water used to house the fish was placed in an enclosed arena with a video camcorder mounted above the tank. Half of the tank was white (light zone), and the other half was black (dark zone). We placed fish individually in this LDT and video-recorded from above for 5 minutes. We quantified duration in the light zone, duration frozen, and number of crosses between zones using BORIS (Friard & Gamba, 2016). The light-dark test is a validated assay to measure stress and anxiety-like behaviors in fish (Gebauer et al., 2011; Maximino et al., 2010). All experimental procedures were approved by the University of Nebraska at Omaha’s Institutional Animal Care and Use Committee (17-070-09-FC).

### 2.3 Statistical Analysis

We performed all statistical analyses using IBM SPSS Statistics (version 29). Using a generalized linear model we tested the main effects of stress coping style (proactive, reactive) and treatment (NACA exposure, control), and stress coping style by treatment interaction effect on the three discrete behaviors (duration in the light zone, duration frozen, and number of crosses between zones). Sex was originally included in the model but later removed after determining there were no significant main effects of sex on all the dependent variables. We applied a Benjamini-Hochberg correction when investigating simple main effects within interaction effects (Benjamini et al., 2001; Storey, 2002). We also performed a principal components analysis (PCA) on the three discrete behavioral measures to assess if variables can be reduced to a composite stress behavior axis. We used the resultant PCA scores as a dependent variable in a generalized linear model to assess whether the stress coping style, treatment, and their interaction had significant effects. When examining main effects of treatment in our generalized linear models, we used one-tail p-values as several studies show that NACA and NAC treatment has stress-reducing effects in zebrafish (Mocelin et al., 2015; Mocelin et al., 2018; Mocelin et al., 2019; Reis et al., 2020).

## 3. Results

Analysis of duration in the light zone showed a significant main effect of stress coping style (*Wald χ*^*2*^ = 22.98, *p* = 1.6 * 10^-6^), where proactive fish spent significantly more time in the light zone than reactive fish (Figure 1A). There was a trend for NACA treated fish to increase time spent in the light (*Wald χ*^*2*^ = 2.62, *p*_*one-tail*_ = 0.053). There was no significant coping style by treatment interaction effect (*Wald χ*^*2*^ = 1.26, *p* = 0.262).

**Figure 1.**
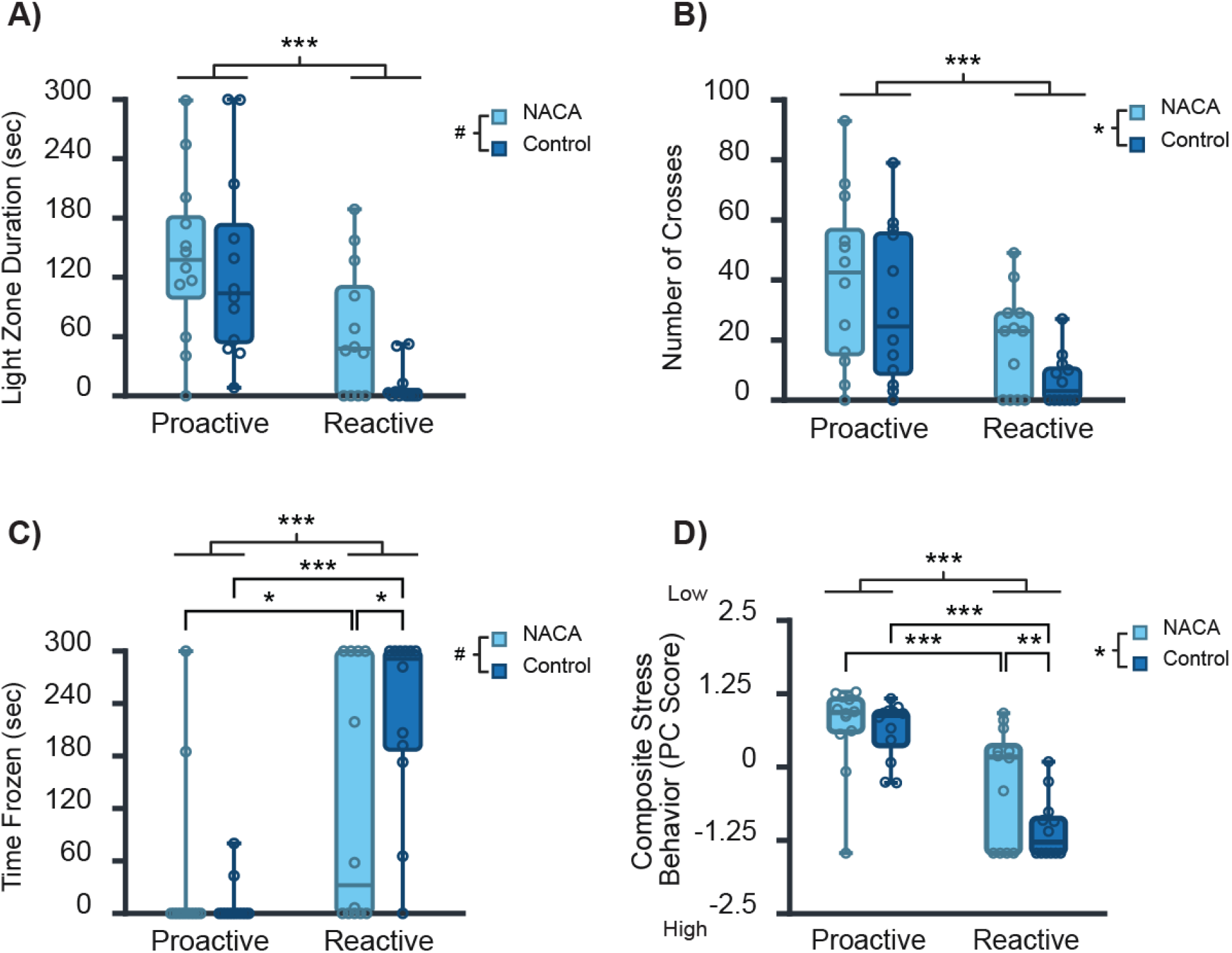
Effect of NACA treatment on discrete and composite stress behaviors for each stress coping style. The (A) duration in the light zone, (B) number of crosses between zones, (C) time frozen, and (D) composite stress behavior score (principal component score) were measured for each fish in the light-dark test. Light blue represents NACA treated individuals and Dark blue represents control (water-treated) individuals. #, p < 0.1; *, p < 0.05; **, p < 0.01; ***, p < 0.001

There was a significant main effect of stress coping style on number of crosses (*Wald χ*^*2*^ = 14.279, *p* = 1.5 * 10^-4^), where proactive fish crossed between zones more frequently than reactive fish (Figure 1B). There was a significant main effect of treatment (*Wald χ*^*2*^ = 3.15, *p*_*one-tail*_ = 0.038) on number of crosses where individuals treated with NACA crossed between zones more frequently than controls. There was no significant coping style by treatment interaction effect (*Wald χ*^*2*^ = 0.1, *p* = 0.756).

There was a significant main effect of stress coping style on freezing duration (Figure 1C, *Wald χ*^*2*^ = 28.28, *p* = 1.05 * 10^-7^), where reactive fish spent more time frozen than proactive fish (Figure 1C). There was a trend for NACA treated fish to have lower amounts of time frozen than controls (*Wald χ*^*2*^ = 1.67, *p*_*one-tail*_ = 0.098). We observed a significant stress coping style by treatment interaction effect on time frozen (*Wald χ*^*2*^ = 5.59, *p* = 0.018). More specifically, NACA treated individuals of the reactive stress coping style significantly decreased their time frozen relative to controls (p = 0.01) but this pattern was not observed between NACA treated or control proactive individuals (p = 0.449). In both control and NACA treated animals, the reactive fish spent significantly more time frozen than proactive fish (p = 5.6 * 10^-8^ and p = 0.037, respectively).

PCA analysis revealed that the three discrete behaviors all loaded onto a single component (composite measure of stress behavior) accounting for 65.7% of the total variance. Time in the light zone and number of crossings loaded positively, whereas time frozen loaded negatively. Resulting PC scores show that more positive scores indicate lower stress-related behaviors. Proactive fish had significantly higher PC scores than reactive fish (*Wald χ*^*2*^ = 43.33, *p* = 4.62 * 10^-11^) (Figure 1D). NACA treated fish had significantly higher PC scores than controls (*Wald χ*^*2*^ = 4.79, *p*_*one-tail*_ = 0.029). We also found a trend for coping style by treatment interaction effect on PC scores (*Wald χ*^*2*^ = 3.46, *p* = 0.063). More specifically, NACA treated individuals of the reactive stress coping style had significantly higher PC scores relative to controls (p = 0.004) but there was no significant difference in PC scores between treated and control proactive individuals (p = 0.817). In both control and NACA treated animals, the proactive fish had significantly higher PC scores than reactive fish (p = 2.36 * 10^-9^ and p = 0.001, respectively).

## 4. Discussion

In response to stressors, individuals will vary in behavioral and physiological responses. While oxidative stress and antioxidants can directly modulate glucocorticoid levels and its effects, the role of individual differences on this relationship has not been well-studied. Notably, non-stressed proactive and reactive zebrafish differ in whole brain levels of biochemical and gene expression markers of oxidative stress and antioxidants (Sunday-Jimmy et al., 2026; Wong & Godwin, 2015), suggesting there may be differing capacities to withstand oxidative insults between the stress coping styles. Our results show that the magnitude of behavioral stress reducing effects of the N-acetylcysteine amide antioxidant differs by stress coping style.

NACA treatment resulted in an increase in duration of time in the light zone, reduced duration frozen, increased number of crosses between zones, and higher PC scores (Figure 1). As adult zebrafish display elevated scototaxis when stressed (Gebauer et al., 2011; Maximino et al., 2010; Wong et al., 2012), increased preferences for the light zone suggest reduced stress levels. Our results are consistent with prior studies in zebrafish and other animals that show reduced stress and anxiety-like behaviors upon exposure to NACA or NAC (Dean et al., 2011; Marcon et al., 2019; Mocelin et al., 2015; Mocelin et al., 2018; Mocelin et al., 2019; Reis et al., 2020; Santos et al., 2017; Shariatmadari et al., 2026). While other stress-altering compounds (e.g., ethanol, fluoxetine, caffeine) have demonstrated zebrafish strain-specific effects, congruency of NACA’s effect on stress behavior between the current study and a prior study using wild-type zebrafish strain suggest differences between strains may be minimal (Dlugos & Rabin, 2003; Klucas & Wong, 2026; Maximino et al., 2013; Pannia et al., 2014; Rosa et al., 2018; Wong et al., 2013). However, as we used a different NACA dose and treatment length to obtain similar stress-reducing effects as a prior study (Reis et al., 2020), it suggests there are individual differences in NACA pharmacodynamics in zebrafish.

Notably, we observed stress coping style by treatment interaction effects on duration of time frozen and the composite stress behavior score. Specifically, NACA’s stress-reducing effects were only observed in individuals with the reactive stress coping style (Fig. 1C & 1D). Prior studies have shown stress coping style specific baseline and neuroendocrine stress-induced oxidative stress and antioxidant profiles (Arnold et al., 2014). Across birds, mice, and fish, those with a reactive-like coping style had elevated plasma, serum, muscle, or brain antioxidant biomarker levels at baseline or in response to an acute neuroendocrine-stressor (Arnold et al., 2014; Costantini et al., 2008; Herborn et al., 2011; Rodel et al., 2022; Sunday-Jimmy et al., 2026). However, the interaction between stress coping style, neuroendocrine stress, and oxidative and antioxidant profiles is complex and varies by context and tissue source (Arnold et al., 2014; Herborn et al., 2011; Pamplona & Costantini, 2011). To our knowledge the current study is the first to show that presumably increasing antioxidant levels (glutathione via NACA) reduces behavioral stress levels in a stress coping style-specific manner. Intriguingly, NACA treatment in reactive fish promoted them to be more proactive-like in their behavioral stress levels; NACA-treated reactive fish did not significantly differ from untreated proactive fish in composite stress scores or duration of time frozen. We speculate that NACA-induced increase in antioxidant capacity may be important for facilitating proactive behavioral response to stress. An alternative explanation could be that display of the reactive stress coping style’s behavioral response to stress is through presumed lower threshold for disrupting redox homeostasis due to lower NACA-mediated antioxidant levels.

The behavioral effects of NACA observed in the current study are hypothesized to arise from its combined enhancement of antioxidant capacity and modulation of glutamatergic signaling. While the current study did not quantify redox biomarker levels or glutamate signaling, multiple studies demonstrate that administering NAC or NACA in zebrafish and other animals increase antioxidant levels and alter glutamatergic signaling in the brain (Bradlow et al., 2022; He et al., 2020; Pedre et al., 2021; Raghu et al., 2021; Reis et al., 2020; Spilere et al., 2025). Inside the body NACA has enhanced ability to cross the blood-brain barrier and is rapidly converted to N-acetylcysteine (NAC) (He et al., 2020; Reis et al., 2020). While NAC can directly act as an antioxidant, it is thought to predominantly serve as a precursor for glutathione antioxidant system (Bradlow et al., 2022; Pedre et al., 2021; Raghu et al., 2021; Spilere et al., 2025). Studies also showed that NAC can modulate a major brain neurotransmitter signaling system for stress and anxiety, glutamate, through the astrocytic cystine-glutamate transporter (Dean et al., 2011; Raghu et al., 2021; Reis et al., 2020). By reducing oxidative stress and influencing cystine–glutamate exchange, we speculate that NACA may dampen excitatory neural activity within the neuroendocrine stress axis (hypothalamic-pituitary-adrenal/interrenal axis. We also hypothesize that glutathione buffering capacity is a key redox mechanism facilitating biasing stress responses to be proactive- or reactive-like. Future studies should quantify brain glutathione and glutamate levels after NACA administration.

In summary, this study demonstrated there are individual differences in the stress-reducing effects of NACA in zebrafish. NACA’s effect was only seen in individuals with the reactive stress coping style and promoted more proactive-like responses within these individuals. These findings underscore the importance of considering individual differences in stress coping styles and redox states when investigating mechanisms of stress and anxiety-like behaviors.

Given NACA’s role in ultimately enhancing glutathione redox signaling, we hypothesize that the reactivity of the glutathione system is key to biasing behavioral and physiological responses to neuroendocrine stressors characteristic of a stress coping style.

## Acknowledgements

We thank UNO’s Animal Care and Use Program staff for zebrafish husbandry. We are grateful to Jamie Corcoran, Princess Sunday-Jimmy, and Tai Prauner for helpful discussions. This project was funded by the National Institutes of Health (5P20GM103427) to RYW and University of Nebraska at Omaha’s Fund for Undergraduate Scholarly Experience to BKS.

## Author Contributions

RYW: Conceptualization, formal analysis, funding acquisition, supervision, visualization, writing – original draft, writing – review and editing

BKS: Conceptualization, funding acquisition, data curation, investigation, methodology

CRG: Formal analysis, writing – original draft

PDD: Conceptualization, funding acquisition, writing – review and editing

